# What should a poor mother do? Influence of host plant quality on oviposition strategy and behavior in a polyphagous moth

**DOI:** 10.1101/2021.06.03.446956

**Authors:** Kristina Karlsson Green, Benjamin Houot, Peter Anderson

## Abstract

To maximise fitness, individuals may apply different reproductive strategies. Such strategies could be phenotypically plastic and vary depending on the environment. For example, when resources are limited females often face a trade-off between investing in offspring quantity and quality, and how she balances this trade-off may depend on the environment. For phytophagous insects, and especially generalist insects, variation in host plant quality could have large effects on mating, reproduction and offspring performance. Here, we study if the polyphagous moth *Spodoptera littoralis*, which selects host plants through experience-based preference induction, also has a flexible allocation between egg weight and egg number as well as in temporal egg-laying behavior depending on larval host plant species. We found that *S. littoralis* has a canalized egg size and that an increased reproductive investment is made in egg quantity rather than egg quality. This increased investment depends on larval host plant species, probably reflecting parental condition. The constant egg weight may be due to physiological limitations or to limited possibilities to increase fitness through larger offspring size. We furthermore found that differences in onset of egg-laying is mainly due to differences in mating propensity between individuals raised on different host plant species. Thus, females do not seem to make a strategic reproductive investment in challenging environments. Instead, the low-quality host plant induces less and later reproduction, which could have consequences for population dynamics in the field.

## Introduction

Reproduction is crucial for individual fitness but it is also a costly engagement that requires large resources. How individuals invest in reproduction could thus be shaped by trade-offs due to resource limitations (Chippindale et al. 1993). To maximize fitness, individuals commonly apply different reproductive strategies (Gross 1996), which include various behavioural, physiological and morphological traits that influence mating and reproduction. Such strategies could either be genetically fixed, such as in the side-blotched lizard *Uta stansburiana* where different genetic colour morphs invest in either offspring quality or quantity (Sinervo et al. 2000), or vary depending on the environment and thus be phenotypically plastic. For example, the social environment could influence mating propensity in the fresh-water isopod *Asellus aquaticus* (Karlsson et al. 2010) and experience of acoustic signals could affect male investment in reproductive organs in crickets (Bailey et al. 2010).

Plasticity is often favourable when the environment varies (West-Eberhard 2003). Plasticity in reproductive strategies can be complex, as this could be induced in the juvenile stage but not expressed until adulthood and in addition, the plastic expression could have consequences for offspring and thus have effects across generations. For example, a plastic expression of reproductive strategies could be affected by the environment that the reproducing individual has experienced previously, e.g. resource acquisition before the reproductive event (Katsuki et al. 2012). In addition, the plastic response could be dependent on the environment that the individual is exposed to during the current mating and reproduction, e.g. characters of the mating partner (Pizzari et al. 2003). The plastic expression could furthermore be either an involuntary consequence of the individual’s condition, for example if individuals in good condition may invest more in mating (Duplouy et al. 2018), or a strategic investment to improve offspring fitness depending on assessment of the particular environment, e.g. sex-ratio allocation of offspring based on perceived host quality (Pexton & Mayhew 2005), and thus being adaptive. A plastic reproductive strategy is expressed in the reproducing adults, and is thus a case of within-generational plasticity, but the strategy could have trans-generational consequences if the strategy modifies offspring phenotypes (Bonduriansky & Crean 2018). One example is when females invest either in larger or smaller eggs, which could have consequences for offspring development and survival Cahenzeli & Erhardt 2013). However, although plasticity may be favorable for adjusting to environmental variation, canalization of traits often occurs in nature, for example when plasticity carries a cost or when the benefits of plasticity are limited (Auld et al. 2010; DeWitt et al. 1998). Thus, individuals may not be able to apply plastic strategies in all possible aspects of reproduction.

For phytophagous insects, the host plant is often of great importance both for mating and for offspring performance and survival (Schoonhoven et al 2005). Host plants commonly vary in quality, both within and between plant species, and females therefore usually select a suitable host plant for their eggs with large care. The host plant species and quality thus have large potential to influence reproductive strategies in insects (reviewed in Awmack & Leather 2002; Moreau et al 2017). For example, female condition is in general influenced by the host plant she developed upon as a larva and the quality of the larval host plant can therefore have a direct effect on the resources available for reproduction, especially when egg production is dependent on nutrients accumulated in the larval stage (Wheeler 1996). Female host plant experience could, however, also influence how she anticipates the environment for her offspring, and she may accordingly adjust her reproductive strategy to maximise their fitness (Cahenzli et al. 2015). Larval host plant quality thus has the potential to influence female reproductive strategies and trade-offs that are governed by resource variation.

Generalist insect species utilize a wide range of plant species, that may come from very different families and thus represent a large spatial and temporal variation in resource quality. Due to this environmental variation, generalists may not be as well-adapted to each of their possible host plant species as specialist insects are to their few host plant species (Rothwell & Holeski 2019; Schapers et al. 2016). It has therefore been proposed that experience-based plasticity would be important for generalist species to manage the variation that multiple host plant species presents them with, for example during host plant selection (Bernays 2001). The reproductive strategies that ovipositing females could apply may, however, consist of several different components other than the actual host plant choice. For example, females across species groups are commonly expected to face a tradeoff between investing in offspring quantity or offspring quality (e.g. weight or size) (Smith & Fretwell 1974, Lim et al 2014). This is also seen in phytophagous insects where females could adjust their egg investment depending on host plant quality (Fox et al. 1997; reviewed in Fox & Czesak 2000). Females may moreover modify their temporal oviposition behavior depending on the environment by adjusting the length or onset of their egg-laying period (Berkvens et al. 2008; Saastamoinen & Hanski 2008). Thus, even if plasticity due to host plant experience is beneficial to generalist insects, it is not known if such plasticity is operating on all or only a selection of the traits.

In the current paper, we aimed to investigate the effects of larval host plant species on reproductive strategies in the generalist moth *Spodoptera littoralis*. This species feeds on a large number of plant species from many different plant families that are of varying quality for the insect and *S. littoralis* exhibits plastic responses in both preference and performance depending on larval host plant species. For example, larval immune function (Karlsson Green In press), performance and adult lifespan differ depending on larval host plant species (Karlsson Green et al unpubl.) indicating important effects of plant species on individuals’ condition. The larval host plant species of parents furthermore have transgenerational effects on their offspring performance (Rösvik et al. 2020). Host plant induced plasticity does, however, not only occur on performance but also on preference in *S. littoralis*. Adults of both sexes have an innate preference hierarchy among host plant species, which can be altered depending on the plant species that they experienced as larvae (Anderson et al. 2013; Lhomme et al. 2018; Proffit et al. 2015; Thöming et al. 2013; Zakir et al. 2017). Thus, one component of the females’ reproductive strategy, host plant selection for mating and oviposition, is plastic and depends on the larval host plant species. Whether the plastic host plant choice is further combined with a flexible oviposition strategy depending on larval host plant species is, however, not known.

Here, we thus address if ovipositing *S. littoralis* females show plasticity in their egg-laying strategy depending on larval host plant species. We use three host plant species that vary in quality as larval food and hypothesize that the differences in host plant quality could induce plastic responses and change the oviposition strategy depending on female host plant experience. The plastic response could however be either a direct consequence of the female’s resource availability during the larval stage, or an adaptive allocation depending on expectations of her own reproductive potential and future offspring environment. In our experiments, we address if females alter investment between egg quantity (number) and egg quality (measured as weight). In addition, we address if females alter their temporal egg-laying behavior and if this is dependent on delayed onset of oviposition or delayed mating. We hypothesise that if the plastic response is a carry-over effect of female condition, females of the most challenging host plant species would have smaller and fewer eggs as well as a shorter and later egg-laying period. However, if the plastic response is an adaptive strategy to compensate for a resource-poor environment, we expect females from the challenging host plant species to invest more in egg quality than in quantity and also to oviposit earlier.

## Materials and methods

### Study species

*Spodoptera littoralis* is a polyphagous and nocturnal moth that feeds on more than 80 different plant species that comes from a wide range of plant families (CABI 2019). The species is a significant crop pest that is present throughout Africa, the Middle East and Southern Europe (CABI 2019). A lab colony of field-collected *S. littoralis* from Egypt is reared at SLU, Alnarp where the animals are raised in climate chambers with controlled settings of 16:8 L:D, 25°C, 60% RH. In all bioassays described below, larvae were reared in groups in plastic boxes (H*W*L 6.5*18*22 cm), feeding detached leaves *ad libitum* until pupation. At the pupal stage, males and females were separated until eclosion and adults were mated at the age of two days. All bioassays were performed in the rearing conditions (16:8 L:D, 25°C, 60% RH).

Cotton (*Gossypium hirsutum*), cabbage (*Brassica oleracea* v. *capitata*) and maize (*Zea mays*), that were used as host plants in the current study, were cultivated from seeds in a greenhouse with controlled settings (16:8 L:D, 25°C, 70% RH). All these species are present in the agroecosystem Egypt where the lab population originates from. Even though all plants are domesticated and that they have different geographic origin, wild related plants to these three crops naturally occur within the distribution of the studied population of *S. littoralis*. This indicates that the evolutionary relationship between the plants and the insect is longer that when cultivation of crops was intensified in this region. The Egyptian population of *S. littoralis* has an innate preference hierarchy in which it prefers cotton and maize over cabbage but this preference hierarchy may shift due to larval induced preference (Anderson et al. 2013; Thöming et al. 2013) which is mediated by olfactory cues (Lhomme et al. 2018). The preference hierarchy is not associated with larval performance (Karlsson Green et al unpubl) as individuals in general have a fast development and large pupal weight on cabbage, which they don’t prefer, but a very poor development on maize, which they prefer over cabbage (Roy et al. 2016).

### Experiment 1: egg investment and egg-laying behaviour

To assess if females alter their oviposition strategy depending on larval host plant species we studied their investment in egg quality vs. egg quantity as well as their temporal egg-laying behavior during the entire life-time of females reared on either cotton, cabbage or maize plants as above. First, a male and a female were introduced into a cylindric mating cage (height 15 cm, Ø 11 cm) provided with honey-water to feed on. A tracing paper was included around the cage walls to oviposit on but no host plant material. To characterize the egg-laying behaviour, we measured the weight of the egg batches every day until the death of the female. We also noted the first day of oviposition and the total number of egg-laying days for each female. To record the number of eggs for the first egg batch, this batch was deposited on filter paper (Whatman GradeNo; 1, Ø 90 mm) inside a glass petri dish (Ø 90 mm) with 1 ml of methanol overnight. The egg batches were photographed and analysed with the ImageJ software. The investment in individual egg weight (i.e. egg size) for each female was then calculated as the total weight of the first egg batch divided with the number of eggs in that batch (number of clutches analysed per treatment: 10≤N≤23).

### Experiment 2: mating propensity

To disentangle if onset of egg-laying behaviour is affected by differences in mating propensity (i.e. time until mating occurs) or differences in the time it takes for the fertilised eggs to develop until oviposition, we performed a mating experiment with individuals reared on either cotton, cabbage or maize. Larvae were reared in groups on detached leaves of either of the three host plants as described above. Two-days old adults that had fed the same host plant diet were put in cylindric mating cages (height 15 cm, Ø 11 cm), one male and one female in each cage, provided with paper to oviposit on and water. No honey was added to the water in this experiment to ensure that differences between treatments were due to larval acquired resources. During the first day of the experiment, the cages were monitored every 45 minutes, for eight hours, to observe if mating occurred or not. The following days, the cages were monitored once every day to record if and when the first egg batch appeared. The experiment was ended when a clutch had been laid or when the female was found dead. The mating experiment was performed in a climate chamber with the same settings as the rearing chamber (16:8 L:D, 25°C, 60% RH). In the experiment, we used a total of 50 pairs (17 reared on cotton, 18 reared on cabbage and 15 reared on maize).

### Statistical analyses

For Experiment 1, the effect of larval host plant diet on egg-laying parameters was analysed using XLSTAT 2012 software (Addinsoft, XLSTAT 2012). The impact of larval host plant species on individual egg weight, the number of eggs in the first batch, total egg weight, onset of egg-laying-and length of the egg-laying period (number of days) was assayed with Kruskal-Wallis tests completed by Dunn’s procedure to obtain multiple pairwise comparisons (at level p = 0.05). An ANCOVA was performed in JMP version Pro 15 to analyse the differences in weight of the first egg clutch depending on host plant species, the number of eggs in the clutch, and their interaction.

To assess differences in mating propensity in Experiment 2, we performed a generalised linear model with binary response variable and logit link-function in JMP version Pro 14. Response variable was whether the pair mated the first day or not and explanatory factor was larval host plant species. To address if a difference in time until the first oviposition event was due to differences in mating propensity or in the time between mating and oviposition, we analysed the number of days between mating and egg-laying for the pairs that we had observed mating to occur with Kruskal-Wallis test. Also in this model, larval host plant was included as the explanatory factor and a total of 36 pairs were analysed of the initial 50 pairs in the experiment (N cotton = 16, N cabbage = 14, N maize = 6).

## Results

In Experiment 1, larval host plant was found to affect egg quantity of the first clutch oviposited, as females reared on cotton laid both a higher number of eggs than females fed on maize (mean eggs ± SD: cotton: 350 ± 122, cabbage: 225 ± 149, maize: 131 ± 55; df = 2, χ^2^ = 18.975, p < 0.0001) and a larger clutch weight (mean mg ± SD: cotton: 20 ± 7, cabbage: 13 ± 8, maize: 7 ± 3, df = 2, χ^2^ = 22.957, p < 0.0001). The ANCOVA revealed that the weight of the first egg clutch was only dependent on the number of eggs in the clutch (F_1,50_ = 327.763, p < 0.0001) and not on host plant species (F_2,50_ = 1.468, p = 0.241) or the interaction between species and egg number (F_2,50_ = 0.819 p = 0.448). Moreover, there were no differences in individual egg weight in the first clutch between the three host plant diets (Fig. 1; df = 2, χ^2^ = 2.476, p = 0.290). We also found that the total egg weight that a female deposited during her lifetime differed depending on larval diet, where females raised on maize had a lower total egg weight than females reared on cotton and cabbage (Fig. 2a; df = 2, χ^2^ = 12.326, p = 0.0002). Onset of egg-laying differed depending on larval host plants as cotton raised females laid there first clutch earlier than cabbage fed females and maize fed females initiated their egg-laying latest of all (Fig 2b; df = 2, χ^2^ = 19.240, p < 0.0001). However, there was no difference in length of egg-laying period depending on larval host plant (Fig 2c; df = 2, χ^2^ = 5.490, p = 0.064).

**Fig. 1.**
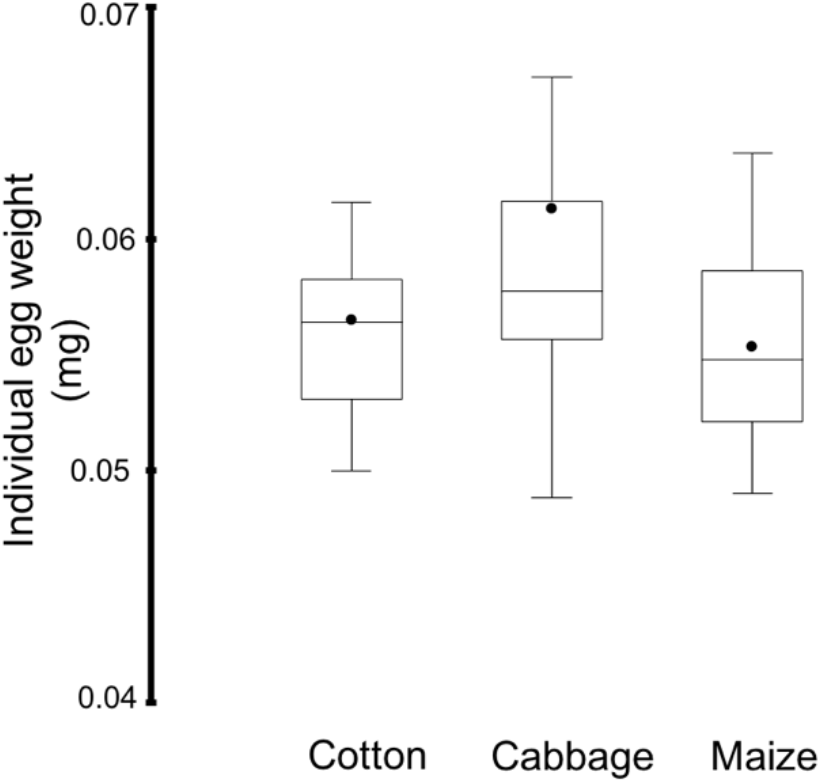
The investment in individual egg weight depending on larval diet in *S. littoralis*. The individual egg weight in the first clutch for females feeding cotton, cabbage or maize, which showed no significant difference (Kruskal-Wallis test with Dunn procedure, p = 0.29). Boxes represents 25^th^ and 75^th^ percentiles and error bars represents the 10^th^ and 90^th^ percentiles. Horizontal lines within boxes represent median value and black dots represent the mean.

**Fig. 2.**
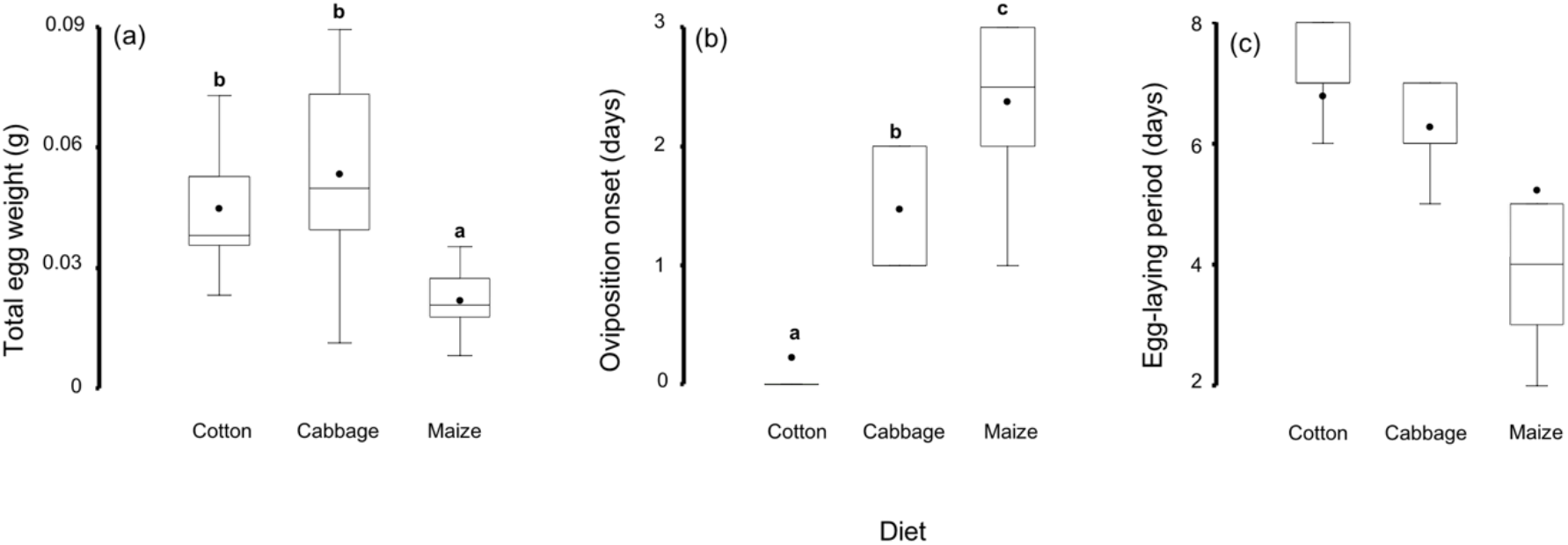
Egg production and temporal egg laying behaviour depending on larval host plant in *S. littoralis*. (a) The difference in total egg weight that females raised on cotton, cabbage or maize deposited during the experiment (p = 0.0002). (b) The difference in oviposition onset (number of days from experiment start until the first egg clutch) for females raised on cotton, cabbage or maize (p < 0.0001). (c) The length of the total egg-laying period for females raised on cotton, cabbage or maize (no significant difference, p = 0.064). Different letters above boxes indicate significant differences at level p = 0.005 in Kruskal-Wallis test with Dunn’s procedure. Boxes represents 25^th^ and 75^th^ percentiles and error bars represents the 10^th^ and 90^th^ percentiles. Horizontal lines within boxes represent median value and black dots represent the mean.

In Experiment 2, we furthermore found that the delay in egg-laying between females reared on different host plants depended on mating propensity, where a higher proportion of pairs reared on cabbage and cotton mated during the first day, than pairs reared on maize (df = 2, χ^2^ = 9.511, p = 0.009, Fig. 3a). There was however, no difference in time between mating and egg-laying between pairs raised on different host plants (df = 2, χ^2^ = 1.113, p = 0.573, Fig. 3b).

**Fig. 3.**
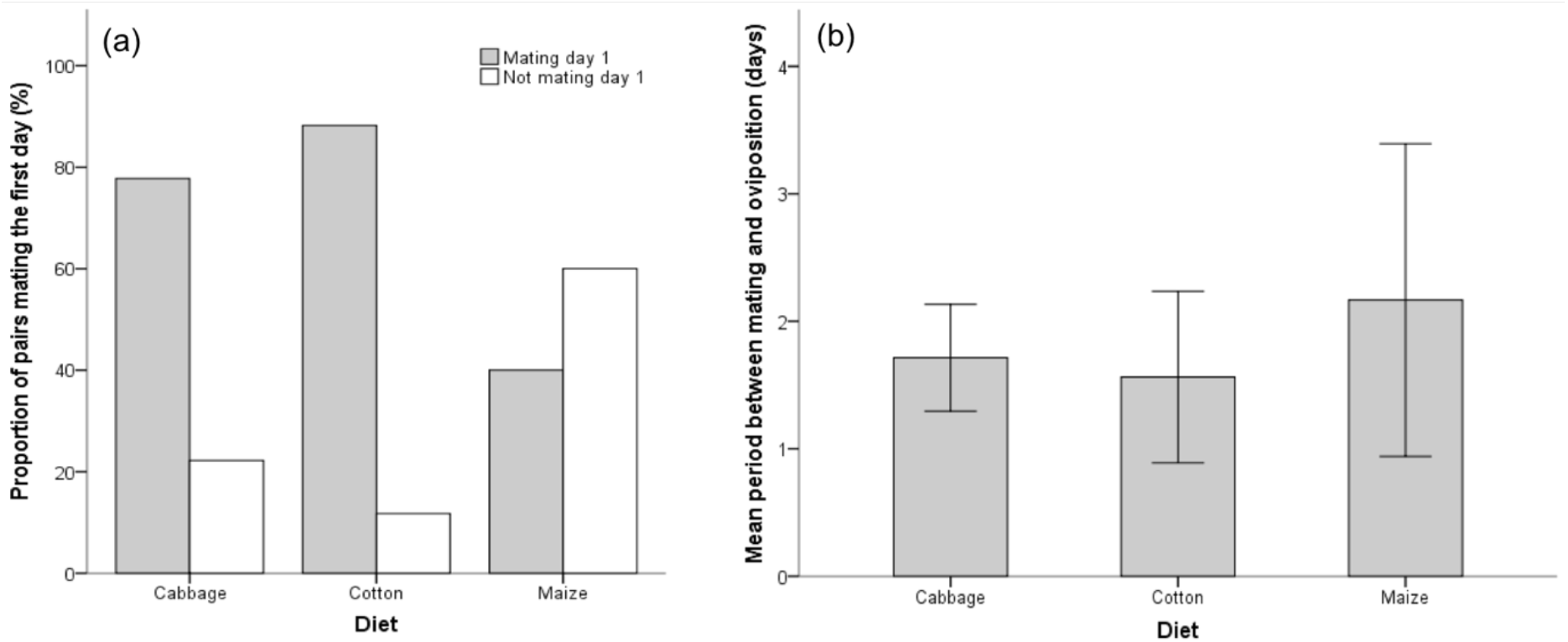
The effects of delayed mating on oviposition in *S. littoralis*. (a) The proportion of pairs raised on different larval host plants that mated during the first day (GLM, p = 0.009). (b) The number of days between mating and oviposition which is equal for all females irrespective of larval host plant species (Kruskal-Wallis test, p = 0.573).

## Discussion

Here, we investigated the potential for larval host plant species to affect reproductive strategies in the generalist and highly plastic moth *S. littoralis*. Our results indicate that larval host plant species has consequences for female reproductive output but that females overall allocate resources to egg quantity rather than egg quality, and thus do not have a plastic investment in egg weight. In addition, the differences in temporal oviposition behaviour may be due to delayed mating for individuals reared on low-quality hosts and thus, both male and female condition may affect the subsequent egg-laying pattern.

A plastic reproductive strategy could be favourable when resources vary in the environment. As female reproduction often is resource limited, a trade-off between egg number and egg weight is often assumed, and females are generally predicted to invest in egg quality in poor environments, given her offspring could then benefit from more resources (Amiri et al. 2020; Cesar and Rossi 2019; Moreau et al. 2017). In our experiments, the lowest quality resource environment for females was maize as this host plant is known to provide poor conditions for larval development which results in low pupal weight (Roy et al 2016; P. Anderson unpubl data). However, as there were no differences in individual egg weight between host plants, our results indicate that females do not adjust the weight of individual eggs. Instead, ovipositing females alter their egg quantity depending on larval host plant and oviposit a larger quantity of eggs when they have developed on a better (high quality) host plant. The allocation strategy is thus likely based on female resource acquisition when her eggs are developing, rather than a flexible decision made in relation to larval host plant quality. In some species, the resources that females have available for egg production is also affected by nuptial gifts and ejaculate size from the males they mated with (South and Lewis 2011; Vahed 1998). The size of such gifts could be dependent both on male genotype and the resources he had access to during his development, i.e. may also be an effect of larval host plant. We currently do not know if nuptial gifts are important in *S. littoralis* but males may produce spermatophores of different sizes (P. Anderson unpubl. data) and as we mated pairs that were raised on the same host plant species, the differences that we found in total egg load between females raised on different plants could also depend on how the larval host plant affects males. In Lepidopteran species, both female and male size has been shown to affect female fecundity (Schapers et al. 2017), however Cahenzli and Erhardt (2013) found that males’ larval resources only had minor effects on egg production.

In general, variation in female size (which may be a result of her larval resource acquisition) within Lepidopteran species has an effect on egg number rather than egg size (Bauerfeind and Fischer 2008), which is consistent with our current results. A lack of flexibility of egg size has also been found in other species (Snell-Rood and Steck 2019) but there is in general little knowledge on the possible physiological factors that may constrain egg size plasticity in insects (Fox and Czesak 2000). Aside of the potential physiological constraints to egg size plasticity, there may be only minor opportunities to increase offspring fitness through egg size and the actual egg size could be a result of selection for maternal fitness rather than offspring fitness, as has been found in Atlantic salmon (Einum and Fleming 2000). There may also be more complex relationships between egg quantity and egg quality in insects than a simple trade off (Fischer et al. 2003).

Rösvik et al. (2020) recently showed indications of transgenerational plasticity on offspring performance in *S. littoralis* depending on parental host plant species during the larval stage. An increased egg investment could be a mechanism behind such transgenerational plasticity (Fischer and Fiedler 2001), i.e. maternal effects, where non-genetic components, such as egg nutrients, are transferred from the mother to her offspring to improve their fitness (Bernardo 1996). However, as the results in our current paper indicate that females do not alter egg size depending on larval host plant species, we suggest that egg size in itself does not explain the mechanism behind the transgenerational effects previously found in *S. littoralis* (Rösvik et al. 2020). Indeed, egg size may not be the only parameter for estimating egg investment and egg quality as the yolk protein content could be unrelated to egg size (Diss et al. 1996). There could therefore still be differences in egg quality due to the composition of the egg content that affects offspring performance. In addition, there may be other pathways for transgenerational effects, such as epigenetics (Berger et al. 2009; Bossdorf et al. 2008; Ho and Burggren 2010) or transfer of microbes (Freitak et al. 2014), that do not alter egg size or weight.

We further found that maize-fed females had a later onset of oviposition in comparison to females fed cotton and cabbage. We interpret from this that females on low-quality hosts do not mate and reproduce at an earlier age in order to increase possibilities of reproduction at a low life-expectancy. Instead, we interpret this pattern as an inability to reproduce rapidly due to poor resource environment they have developed in. The delay in onset of egg-laying that we found for individuals reared on cabbage and maize could be due to either a longer time to mature to mating or for eggs to mature following fertilisation, or both. For cabbage fed-females, our mating experiment showed that they mated as early as cotton-fed females and had a similar time between mating and oviposition, thus indicating a difference between experiments in whether there is a delay in oviposition onset or not. However, for maize-fed females this result was consistent across experiments and, as our mating experiment revealed that both cotton-fed and cabbage-fed females mate earlier than maize-fed females, we suggest that the difference in oviposition pattern for maize-fed females is mainly dependent on a delay in mating.

Mating behaviour and investment often depends on the individual’s condition (Buzatto and Machado 2014; Candolin 1999; Perry and Rowe 2010) and could thus depend on either or both of the sexes. For example, male insects are expected to select females based on her fecundity, i.e. her body size (Bonduriansky 2001); as maize-reared individual of *S. littoralis* in general are small (Roy et al. 2016; P. Anderson unpubl data) a low male interest in these females could be a reason for the delayed mating. Moreover, previous studies on *S. littoralis* have shown that females begin pheromone calling for males earlier on host plants than on non-host plants (and on undamaged plants compared to herbivore-damaged plants) (Sadek and Anderson 2007; Zakir et al. 2017). It is possible that larval host plants of different quality could induce similar temporal differences in calling behaviour. Whether it is one of the sexes or both that mature at a later stage may affect the operational sex ratio in the adult population and thus have consequences for sexual selection and mating behaviour (Karlsson et al. 2010; Moura and Gonzaga 2019). A delayed mating, could moreover affect the reproductive output if older females lay less eggs, as in the Codling Moth, *Cydia pomonella* (Vickers 1997). In addition, a delay in the time needed to reach the reproductive phase could result in increased risk of predation before they are able to produce any offspring.

Irrespective of the causes, the delay in mating and the subsequent later oviposition in maizefed individuals, further amplify the differences in moth performance on these three plants species as the generation time on maize is additionally extended. Populations that inhabit this low-quality host could thus suffer from several negative effects on reproduction that likely have consequences on population growth. Interestingly, despite these negative consequence of maize as a host plant, previous research has shown that *S. littoralis* that individuals that have been reared on maize as larvae prefers maize over other host plant species (e.g. Thöming et al. 2013). Together with our results, which indicate that females do not invest in offspring to make them better suited for a low-quality host, this may be interpreted that reproductive plasticity in females has evolved to improve female fitness and not offspring fitness. However, seemingly negative effects on reproduction at some host plant species could in nature be balanced by differences in exposure to predators and parasitoids if low-quality hosts provides an enemy free space (Murphy and Loewy 2015; Singer et al. 2004). It is thus relevant for both fundamental science and pest management understand how ecology affects female reproductive strategies and which consequences this has for population dynamics.

As shown here, larval host plant species affect some, but not all, aspects of the reproductive strategies in the generalist *S. littoralis*. We interpret our results to be due to female condition and her larval resource acquisition rather than a strategic investment to maximize offspring fitness. However, to fully understand the oviposition behaivour will require further studies on how offspring fitness is altered by female strategies. In this context, it would be valuable to consider both higher trophic interactions and the (co-)evolutionary history of plant species and *S. littoralis*. The lack of egg size investment raises further questions on transgenerational plasticity; if the offspring are not affected by maternal condition through increased energy allocation, what other mechanisms for maternal effects, such as epigenetics or transfer of microbiota, may be more relevant in this system? Finally, research on how host plant species affect female reproductive strategies is not only of importance to understand fundamental aspects of ecology and evolution; how egg-laying behaviour of pest insects differ between host plants may also affect how we predict pest outbreaks and optimise biological control (Moreau et al. 2016).

## Acknowledgements

We are thankful to Elin Isberg, Elisabeth Marling, and Zahra Mouradinour for help with the experiments and insect rearing. We also thank Audrey Bras, Axel Rösvik, Björn Eriksson, Mattias Larsson, Fredrik Schlyter and Paul Egan for providing valuable comments on a previous draft of this manuscript. Funding was provided from the Swedish Research Council (2014-6482) and Marie Sklodowska Curie Action (INCA 2014-6418) to KKG and from Carl Trygger’s Foundation to PA.

